# A hybrid mortality prediction model of patients with cavitary pulmonary carcinomas

**DOI:** 10.1101/341156

**Authors:** Maliazurina Saad

## Abstract

To date, developing a reliable mortality prediction model remains challenging. Although clinical predictors like age, gender and laboratory results are of considerable predictive value, the accuracy often ranges only between 60-80%. In this study, we proposed prediction models built on the basis of clinical covariates with adjustment for additional variables that was radiographically induced. The proposed method exhibited a high degree of prediction accuracy of between 83-92%, as well as overall improvement of between 6-20% in all other metrics, such as ROC Area, False Positive Rates, Recall and Root Mean Square Error. We provide a proof of concept that there is an added value for incorporating the additional variables while predicting 24-month mortality in pulmonary carcinomas patients with cavitary lesions. It is hoped that the findings will be clinically useful to the medical community.

## Introduction

The prediction of reduced-life expectancy or mortality is central to personalized medicine. It is a complex question since the cause of death varies. The underlying status of body undeniably influence mortality, but may not be listed as primary diagnosis. Traditionally, future mortality predictions have been studied through patients’ medical records, such as laboratory test results, vital signs and demographic information from Electronic Medical Records (EMR) ^[^^1^^-^^3^^]^. Recently, radiographic induced markers, which is commonly known as radiomic markers, started to show potential in a prognostic model ^[^^4^^]^. The latter claimed to be at an advantage due to its non-invasive nature (no blood sample or biopsy needed) while improving accuracy prediction of mortality. In this article, we will investigate the performance of prognostic models developed using both markers, hereafter termed as CR for clinical records and RM for radiomic markers, to prove or nullify the finding in ^[^^4^^]^. Our hypothesis is that RM will boost a model performance to that of traditional model that is based on CR data only.

The models were developed as a risk stratification of 24-month mortality for patients with cavitary lesion in pulmonary carcinomas. A cavitated lesion, as shown in Fig. 1, is a relatively common finding in chest imaging and has been associated with a worse prognosis ^[^^5^^-^^6^^]^, hence the motivation behind this work. Pathologically, it is defined as “*a gas-filled space within a zone of pulmonary consolidation, mass, or nodule, produced by the expulsion of a necrotic part of the lesion via the bronchial tree*”. Radiographically, it is referred to as “*a lucent area within the lung that may or may not contain a fluid-level that is surrounded by a wall usually of varied thickness*”^[^^7^^]^. Although the mechanism of cavity formation is often difficult to ascertain, cavitation in lung cancer most often results from rapid tumor growth that exceeds the blood supply with resultant central necrosis. The goal of this study is to classify patients into two groups: those who survived 24 months (alive) and those who are not (dead), according to the given variables. This study could serve as a decision support tool in therapeutic strategy as well as identifying the increased risk of short-term mortality.

**Figure 1.**
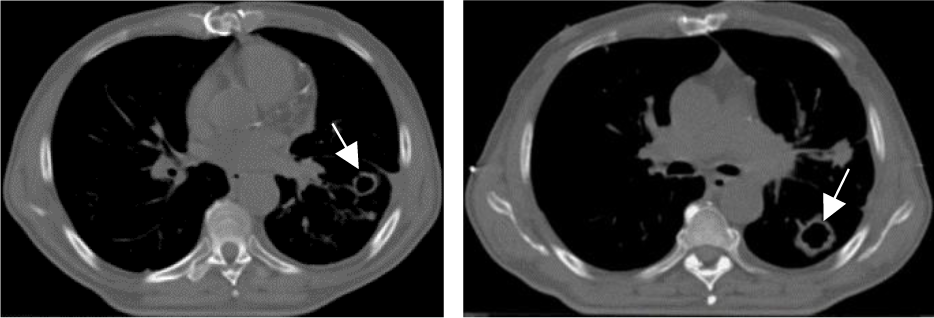
Cavitary lesions in CT imaging

## Materials and Methods

### 2.1 Clinical materials

Imaging and clinical data of patients diagnosed with primary non-small cell lung cancer were obtained from a public repository TCIA ^[^^8^^]^. A total of 47 patients were curated from two cohorts: Radiomics ^[^^9^^-^^11^^]^ and Radio-genomics ^[^^12^^]^. Inclusion criteria encompassed patients with measurable cavitary lesion. Those with more than one cavity were excluded. The acquisition protocol varied slightly for different patients, depending on each patient’s size. Exposure settings ranged from 120 to 140 kVp, with a tube current of 40 to 449 mAs. Pixel spacing and slice thickness were constant at 0.977 mm and 3.0 mm respectively throughout CMS and Siemens vendor, while the rest varied from 1.0 to 5.0 mm and 0.6 to 1.5 mm respectively. The images were reconstructed at 512 x 512 pixel matrices.

### 2.2 Framework description

The first layer in Fig.2 depicts tumor delineation stage as described in^[^^13^^]^. Target lesions were volumetrically delineated using this automatic approach, with an experienced radiologist overseeing and validating the segmented boundaries. The middle layer represents candidate variables included in this study. (A) is a cross sectional view of CT scans, where *N* is the total number of slices., and (B) is a single slice representation, in which *R_x_* and *R_y_* denote the outer and inner radius respectively. *W* indicates the width or wall thickness (WT) of a cavitated tumor. Equation (1) demonstrates the conventional technique of measuring *W.* In contrast, Eq. (2) is a novel way of measuring *W* by taking depth into consideration, hence termed as wall deepness (WD). Note that WD is a 3-dimensional RM compared to WT, which neglected the depth information.
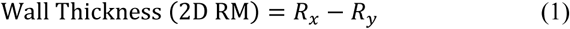

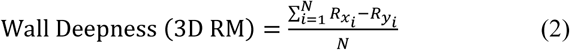

**Figure 2.**
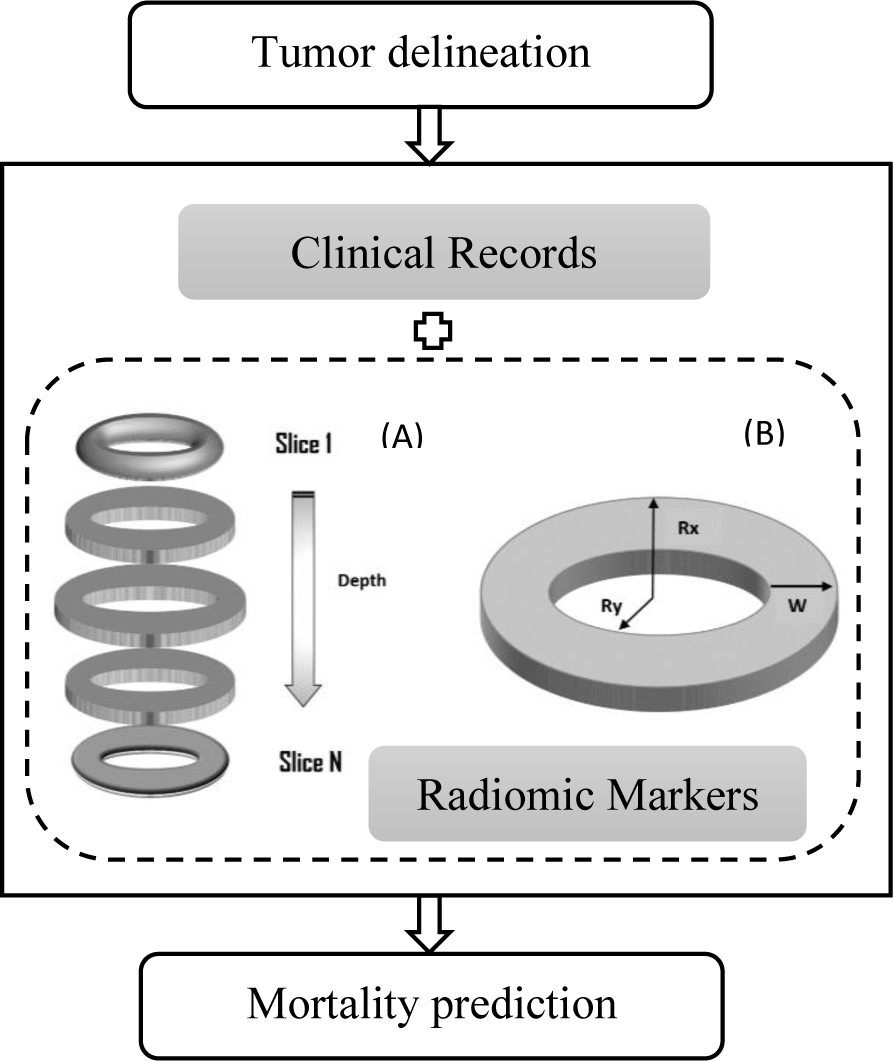
Block diagram of the proposed predictive model

The candidates’ variables *X*={*X*_1_, *X_2_*, … *X*_n_} were designed as shown in Table 1. The final layer is designated for the prediction of vital status (alive or dead) based on the given variables. We designed three test cases based on the input variables to validate our hypothesis as follows:

- CR only
- CR + WT (2D_CR)
- CR + WD (3D_CR)

**Table 1.**
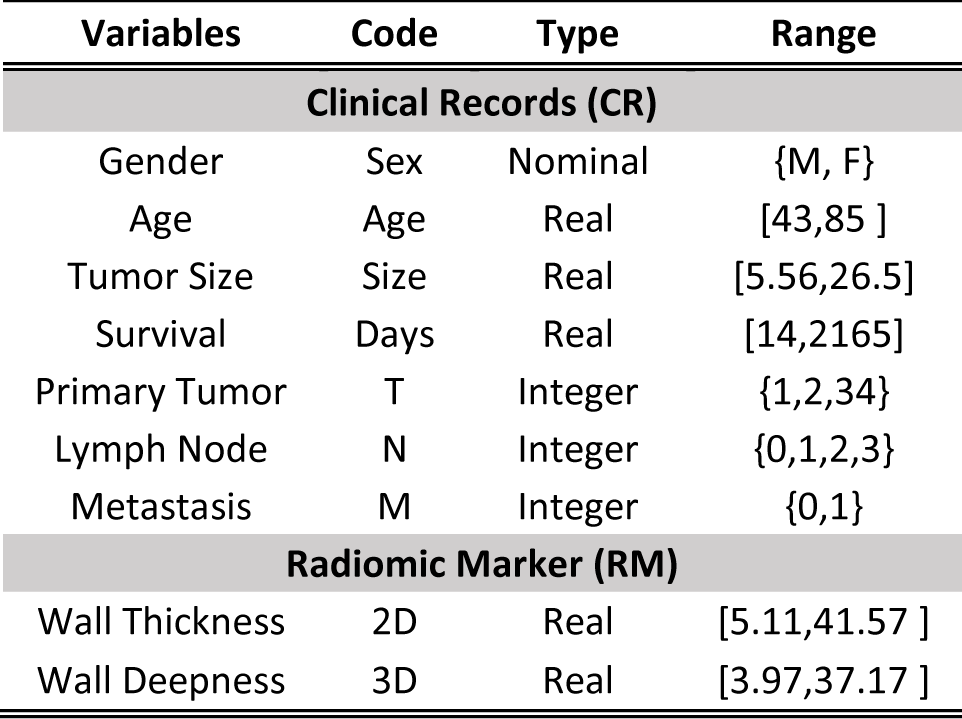
Candidates’ variables under CR group are based on demographics, staging and last day to follow-up data. 2D and 3D wall thickness measurements are included under RM group.

| Variables | Code | Type | Range |
| --- | --- | --- | --- |
| **Clinical Records (CR)** |  |  |  |
| Gender | Sex | Nominal | {M, F} |
| Age | Age | Real | [43,85] |
| Tumor Size | Size | Real | [5.56,26.5] |
| Survival | Days | Real | [14,2165] |
| Primary Tumor | T | Integer | {1,2,34} |
| Lymph Node | N | Integer | {0,1,2,3} |
| Metastasis | M | Integer | {0,1} |
| **Radiomic Marker (RM)** |  |  |  |
| Wall Thickness | 2D | Real | [5.11,41.57] |
| Wall Deepness | 3D | Real | [3.97,37.17] |

Data partitioning was performed through a 10-fold cross validation ^[^^14^^]^. Six off-the-shelf classifiers were carefully selected to run those test cases. Each one of them is detailed as below:

#### Bayesian classifier (Naïve Bayes)

Supposed there are ‘m’ classes {C_1_, C_2_ … C_m_} and let *X* be the predictors {*X*_1_, *X*_2_ …*X*_n_}. The classification is done to derive maximal p (*C_i_*|*X*) as shown in Eq.(3) ^[^^15^^]^
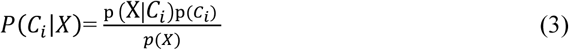

#### Logistic regression

Equation (4) explains how a logistic regression modeled dichotomous dependent variables to describe the relationship with the explanatory variables ^[^^16^^]^
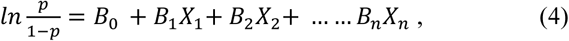

where beta parameters *B*_0_, *B*_1_ … *B*_n_ are modeled by means of maximum likelihood (MAL) and *p* is the probability of presence of the characteristic of interest.

#### Adaptive boosting (Adaboost)

Adaboost combines the output of weak learners into a weighted sum that represents the final output of the boosted classifier, hence also known as ensemble classifier ^[^^17^^]^ as shown in Eq. (5)-(6).
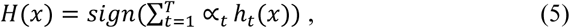

*h_t_*(*x*) is the output of weak learner *t* for input *x,* and ∝*_t_* is the weight assigned to learner *t (0.5*ln(1-E)/E)*, where *E* is an error rate. Once the weak learner trained, the weight of each training sample will be update as in Eq. (6), where *D_t_* denotes weight at previous level, *Z_t_* represents the sum of all weights and (x_i_, y_i_) is a training sample.
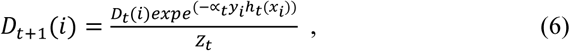

#### Bagging

Another type of ensemble classifier uses the bootstrap aggregating method ^[^^18^^]^ that will reduce the variance of an estimation by averaging together multiple estimates, or in this case the weak learners. Eq. (7) describes this work.
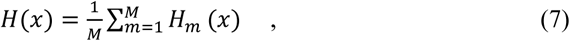

where *H_m_* (m=1 … M) is a sequence of classifiers created by modifying the training samples with a replacement approach.

#### Support Vector Machines

A discriminative classifier that defines an optimal separating hyperplane ^[^^19^^]^. Consider training sample 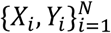: *where X_i_* is the input pattern for i^th^ instance and *Y_i_* is the corresponding target output {-1, 1}. The hyperplane equation is given by
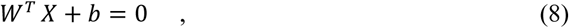

Where *X* denotes an input vector, *W* an adjustable weight vector and *b* is a bias. Thus, the decision rule is defined as
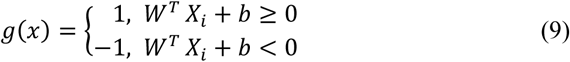

#### Neural Network

A neural network as demonstrated in Fig. 3 was developed. Eight clinical attributes were accepted as the inputs and the training of the network was done with the back propagation algorithm. Eq. (10) explains how neural networks process inputs mathematically
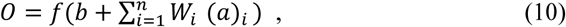

where *0* is the output, (*a*)*_i_* are n inputs, *W_i_* are summation of weights, *b* is the bias and *f* is the activation function ^[^^20^^]^.

**Figure 3.**
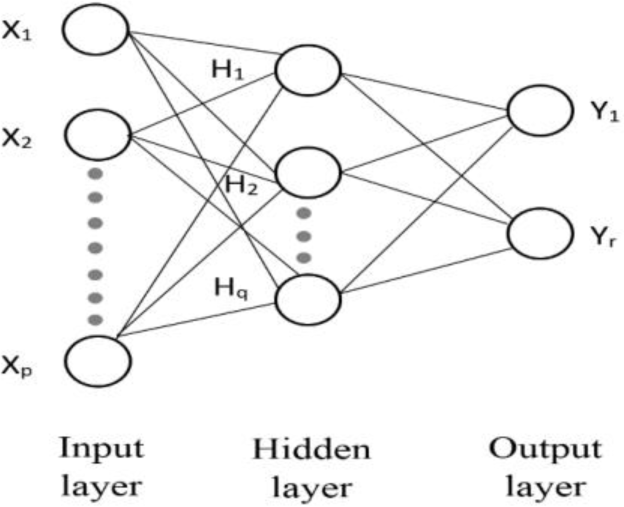
The number of neurons are p=8, q=5 and r=2 for input, hidden, and output layer respectively

## Results

### 3.1 Performance Evaluation

Eighteen models were tested from the combination of six classifiers and three test cases as previously described in section 2.2. The metrics of interest are listed in Eq. (11)-(15).
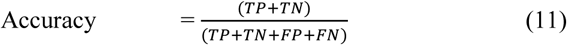

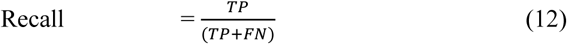

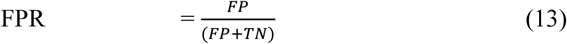

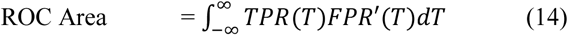

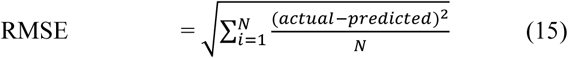

where;

- TP: Number of patients predicted dead that are actually dead
- FP: Number of patients predicted dead that are actually alive
- TN: Number of patients predicted alive that are actually alive
- FN: Number of patients predicted alive that are actually dead
- TPR: True positive rate or recall
- FPR: False positive rate
- T: The integral boundaries of trapezoidal area
- N: Total number of testing subjects
- RMSE: Root mean squared error

### 3.2 Mortality prediction

The performance is evaluated through different version of data fusion: the non-hybrid model (CR only) and two hybrid models (2D_CR and 3D_CR). 2D and 3D refer to the radiomic marker of wall thickness, WT and WD, respectively. All five metrics demonstrated improvements in both hybrid models, with the 3D version to some extent and also outperformed its 2D counterpart. Fig. 4 (a) showed that both hybrid models scored higher **Accurac**y compared to the CR model. The novel 3D marker seemed to improve the prediction slightly higher than the conventional one in all classifiers, except for logistic regression. Similar boosting pattern was recorded by **Recall,** where the sensitivity improved across all the classifiers tested, with considerable differences noticed particularly in the 3D model as depicted by Fig. 4 (b). In contrast, **FPR** in Fig 4. (c) showed significant reduction in all classifiers tested for both hybrid models compared to the CR model. Area under the ROC curve or **ROC Area** reported increasing values in four of the classifiers: Naïve Bayes, Support Vector Machines, Adaboost and Bagging, while remaining basically unchanged in the Neural Network. Interestingly, logistic regression showed mixed performance, where the CR model scored higher than the 3D model but remained lower than the 2D model as demonstrated in Fig. 4 (d). Lastly, the final metrics, **RMSE,** showed substantial reduction in both hybrid models in all classifiers, except logistic regression. The 3D model scored the worst **RMSE** value as shown in Fig. 4 (e). Table 2 summarized the average boosting performance of the hybrid models to that of the traditional CR model. It is noticeable that the 3D model outperformed the 2D model in all cases, except the **ROC Area**.

**Figure 4.**
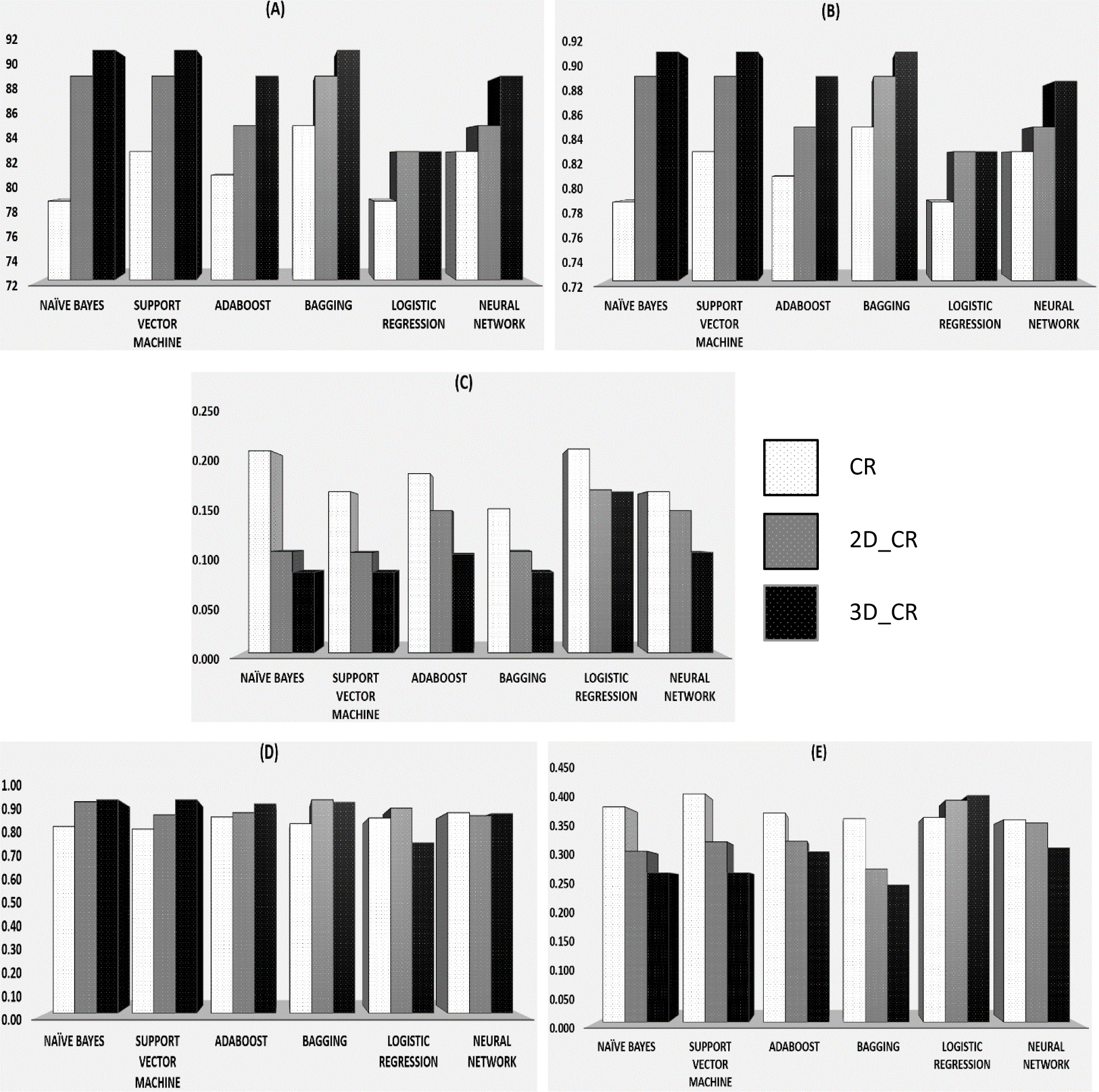
Performance metrics reported by 18 models in terms of A) Accuracy, B) Recall, C) FPR, D) ROC Area and E) RMSE. Note that CR, 2D_CR and 3D_CR represent the three test cases (CR only), (CR+WT) and (CR+WD) respectively.

**Table 2.** Overall performance improvement over the traditional CR model.

| Metrics | 2D_CR (%) | 3D_CR (%) |
| --- | --- | --- |
| Accuracy | 6.6 | 9.6 |
| Recall | 6.6 | 9.5 |
| FPR | 28.2 | 43 |
| ROC Area | 6.5 | 5.9 |
| RMSE | 12.2 | 19.9 |

## Discussion

The experiments demonstrated promising results despite the small dataset and our exclusion criteria that eliminate patients with more than one cavity for the sake of simplicity. We have compared six classifiers on different version of data fusion and noticed that the best results were achieved mostly with the 3D model (3D_CR). Although fusing the 2D marker with the traditional model already improved the prediction ability, we were able to enhance the performance further with the novel 3D marker introduced in this study. These observations confirmed the findings by Carneiro et. al^[^^4^^]^. Primitive classifiers such as Bayesian and Support Vector Machine generally demonstrated major improvement over the traditional model compared to more sophisticated classifier like the neural network, which seemed to be more stable (less significant jump) in terms of performance. We believe this is because of the limited testing subjects that we had. Deep learning classifier is known to be demanding in term of the size of the testing dataset^[^^21^^]^. It is noteworthy to mention that meta-algorithm or ensemble classifiers - Adaboost and Bagging - showed considerable improvement as well. Although they are less significant than the primitive classifiers, they are better than the neural network. Meanwhile, the mixed behaviors of logistic regression merit further investigation. At least two metrics - ROC Area and RMSE - recorded better output in the traditional CR model. We suspected the types of CR variables that mostly fall under nominal or integer as the culprit. This should be an interesting study for the future, provided we have more subjects and more radiomic markers invented.

## Conclusion

In this paper, we showed proof of concept experiments that recommend radiomic markers as potential future prognostic indicator. The findings suggested that radiographically induced marker, if utilized properly, can enhance medical outcomes to some extent. While the experiments were demonstrated using Chest CT imaging, the proposed models could be easily extended to other imaging modalities and patients’ outcomes

